# Divergent ethanol drinking phenotypes are linked to region-specific dysregulation of serotonin systems in the mouse brain

**DOI:** 10.64898/2026.01.30.702832

**Authors:** Brianna E. George, Elena Vidrascu, Sofia Neira, Miles Devine, Thomas L. Kash

## Abstract

Excessive alcohol drinking is a leading cause of preventable death in the United States. High alcohol consumption and persistent drinking despite adverse events, also known as compulsive drinking, are key criteria that contribute to the development and progression of alcohol use disorder (AUD). There is a clear need to better understand the mechanisms that support these related but distinct behaviors. The serotonin (5-HT) system has been associated with alcohol consumption and risk of alcohol dependence, however given the complexity of this system, there remains much to discover regarding specific alcohol related phenotypes. The current study uses a combination of volitional home-cage drinking and operant conditioning to phenotype mice based on ethanol intake and persistence of alcohol drinking following quinine adulteration, a model to study compulsive drinking. Brain tissue of 10 regions known to be implicated in regulating executive function, reward, and stress was collected, and gene expression of serotonergic receptors, transporters, and enzymes was quantified. Three opioid receptors were included given their well-established roles in alcohol-related behaviors and interactions with the 5HT system. Region-specific gene expression patterns emerged, with serotonergic and opioid receptor expression differentially associated with alcohol drinking phenotype. 5-HT and opioid receptors displayed opposing directionality across regions, consistent with functional heterogeneity within the system. These findings identify region-specific molecular alterations following chronic alcohol that may contribute to individual differences in alcohol drinking phenotypes, highlighting candidate targets for biomarkers of increased alcohol use disorder susceptibility or as interventions aimed at preventing the progression of AUD.

## Introduction

High-risk alcohol use remains one of the leading causes of preventable death in the United States with approximately 178,000 deaths each year^1^. While the prevalence of lifetime alcohol use in adults 18 and older is nearly 85%, the prevalence of alcohol use disorder (AUD) in the past year is around 10%^2^. Of those individuals meeting AUD criteria, approximately 59% were classified as having a mild disorder, 20% had a moderate disorder, 20% moderate AUD, and 20% severe AUD, underscoring a need to better understand the neurobiology and behavioral phenotype underlying susceptibility to excessive drinking and AUD.

In clinical practice, the Diagnostic and Statistical Manual of Mental Disorders 5 (DSM-5) is the current framework used to diagnose AUD severity, and is determined by the total number of criteria met in a 12-month period^3^. The 11 AUD diagnostic criteria can be largely classified into two core domains: excessive alcohol use and persistent use despite negative consequences, and increased presence of these criteria is directly linked to worsened severity and functional outcomes. Traditionally, preclinical models of volitional alcohol drinking in rodents employ paradigms that assess alcohol intake and preference using a two-bottle choice (2BC) drinking in the dark (DiD) procedure in the homecage^4–6^. While these binge-like models capture important aspects of alcohol drinking behaviors, such as motivation and reinforcement, they do not assess aversion-resistant drinking, a behavior that has been used to study alcohol consumption despite negative consequences. Additionally, assessment of drinking is temporally constrained to total volume measurements over long periods (30 minutes to several hours), providing limited resolution into critical processes underlying alcohol consumption, such as drinking structure or decision-making^7–10^. Recently, a study established the Structured Tracking of Alcohol Reinforcement (STAR) framework that combines traditional 2BC DiD binge drinking with operant reinforcement procedures to behaviorally phenotype subjects based on multivariate factors, including alcohol intake and aversion-resistant drinking^11^. This model has allowed researchers to delineate individual drinking differences and identify endophenotypes to dissect neurobiological mechanisms contributing to AUD.

The serotonin (5-hydroxytryptamine, 5-HT) system plays a critical role in the regulation of alcohol intake, tolerance, and dependence. Dysregulation of 5-HT neurotransmission has been linked with higher levels of alcohol drinking and an increased susceptibility to dependence^12,13^. However, the relationship between 5-HT neurotransmission and the etiology of alcohol drinking behavior and AUD pathology is not fully understood. Both the clinical and preclinical literature have shown conflicting results regarding the 5-HT system functioning, with reports of increased or decreased functioning being reported across many studies^14^. These mixed findings could be due to diverse function of 5-HT circuitry governing separate motivational or affective aspects of alcohol drinking and the heterogeneous roles of 5-HT receptor subtypes. In the brain, 5-HT neurons are located in the midbrain raphe nuclei, including the dorsal and median raphe nuclei (DRN and MRN), and send extensive projections throughout the brain where they innervate a vast majority of fore- and mid-brain regions. In these downstream regions, 5-HT signals through receptors from one of seven receptor families, designated 5-HT1-7. The 5-HT receptors encompass both G_αi/o_ (5-HT1 and 5-HT5) and G_αq/s_ (5-HT2, 5-HT4, 5-HT6, and 5-HT7) G-protein coupled receptors (GPCRs) or ionotropic receptors (5-HT3). Some of these receptors also function as presynaptic somatodendritic (5-HT1A) or terminal (5-HT1B and 5-HT1D) autoreceptors where they regulate 5-HT release, tone, and activity through negative feedback mechanisms^15^. The multifaceted nature of serotonergic regulation and its overlapping effects on behavior have made functional attribution within the serotonin system challenging. The endogenous opioid system has also been linked to both AUD as well as regulation of 5HT and represents a potential important druggable system that may be targeted synergistically with the 5HT system.

Thus, the current study aimed to better understand the relationship between alcohol drinking phenotype and downstream alteration of serotonin and opioid system machinery and modulators. Using the STAR protocol, mice were phenotyped based on alcohol drinking parameters and then we utilized high-throughput real-time polymerase chain reaction to measure the relative expression of 5-HT and opioid g-coupled protein receptors (GCPRs), transporters, and enzymes across ten brain regions.

## Materials and Methods

### Animals

Adult (>8 weeks) male C57BL/6J (n=24) mice (Jackson Laboratories, Bar Harbor, ME, USA) were allowed to acclimate for at least one week prior to all experiments. Mice were housed in groups of 4 and maintained under a reverse 12/12h light-dark cycle with *ad libitum* water access available via rack-maintained lixit in individually ventilated cages. Prolab Isopro RMH 3000 (LabDiet, St. Louis, MO, USA) was provided daily in measured amounts (3g/animal/day) to maintain healthy adult weight as previously described^11^. A cohort of age-matched males (n=12) were group housed and maintained under the same feeding procedure as alcohol-naïve controls for the qPCR analyses. Two control mice were used for practice tissue collection to optimize procedures and were excluded from all final analyses. All experiments were conducted in accordance with the guidelines of UNC-CH’s Institutional Animal Care and Use Committee (IACUC).

### Operant Procedures

Operant experiments were conducted using a modified version of the STAR protocol^11^. Operant conditioning experiments were conducted in operant conditioning chambers inside sound attenuated cubicles (Med Associates, St. Albans, VT, USA). Each chamber was equipped with two retractable levers located on the back wall with a cue light located above each lever. A retractable sipper connected to a lixit needle valve (Med Associates, ENV-352-2M) was positioned in the center on the front wall of the chamber. Licks were detected and recorded using a contact lickometer (Med Associates, ENV250).. All sessions were conducted during the dark cycle between 0900-1300 and were 1 hour in duration. Alcohol (15% v/v) was diluted from 95% stock in water and was used as the reinforcer. Quinine-adulterated alcohol (250, 500, 750, and 1000 μM) was made daily in 15% alcohol.

### Magazine Training

Mice were placed in the operant chamber with the sipper lixit valve extended continuously and 15% alcohol available freely. Session was terminated and lixits were retracted after 1hr. Mice were retained on magazine training until they reached more than 100 licks in a session. Following acquisition of magazine training, mice were moved to operant conditioning the following session.

### Operant Conditioning

At the start of each session, two levers were extended. One lever was designated the active lever and the other was designated as the inactive lever. Responses on the active lever led to illumination of a cue light above the lever, retraction of that lever, and extension and access to the alcohol lixit for 30 seconds. Responses on the inactive lever had no programmed consequences. Mice were trained on an FR1 reinforcement schedule until they reached a minimum of 10 reinforcers in a single session and then were maintained on FR3 for the remainder of the study.

### Pre- and Post-Binge Operant Testing

The pre- and post-Binge phases were 7 days in length. Days 1-3, 15% alcohol alone was provided during sessions. For days 4-7 increasing concentrations of quinine-adulterated alcohol (250, 500, 750, and 1000 μM in 15% alcohol) were presented with only one concentration provided per day. All sessions were 1hr in length and conducted under FR3, 30sec parameters. All mice were weighed at the end of each session and returned to their group-housed cage at the end of each day.

### Homecage 2 Bottle Choice Testing

The day after the last pre-binge testing session, animals were provided access to alcohol in individual homecages under the common 2 bottle choice drinking in the dark (DiD) procedure. Each animal was placed individually in a clean homecage and given at least 30 minutes to acclimate prior to onset of bottles. Two bottles containing either 15% alcohol (v/v) or water were placed on each cage. Bottle positions were alternated each session to prevent side preferences. All drinking sessions started 3 hours into the dark cycle (1000). Homecage sessions were run 5 days a week with day 1-4 lasting 2hrs in duration and day 5 lasting 4hrs. Bottles were weighed prior to the start and end of each session as well as 2hr into the 4hr sessions. Following the end of each session mice were weighed and returned to their group-housed cage until the beginning of the next session. Binge homecage testing was conducted for a total of 14 days, sequentially with 5 days of drinking and 2 days of abstinence repeating.

### Drinking Phenotypes

Phenotyping was calculated and established following completion of all drinking experiments as previously described^11^. Briefly, each subject’s average lick contacts over the 3 alcohols only sessions were normalized to the mean of licks of all subjects for alcohol only sessions and expressed as a percent. Additionally, each subject’s average lick contacts over the 4 quinine-adulterated alcohol sessions were normalized to the mean of all subjects’ quinine adulterated lick contacted and expressed as a percentage. Subjects were classified as low drinkers if both their percentages of alcohol only and quinine adulterated drinking were less than 100 percent. Subjects were classified as high drinkers if their alcohol only percentage was greater than 100 percent; however, their quinine adulterated percentage was less than 100 percent. The remaining subjects were classified as quinine-resistant and had quinine-adulterated percentages that were higher than 100%.

### Tissue Collection

The day after the final post-binge operant session, mice were anesthetized with isoflurane, decapitated, and brains were rapidly removed and flash frozen in chilled isopentane and stored at - 80ºC until processing. Frozen whole brains were sectioned in a prechilled adult mouse brain matrix and individual slices were collected in 1mm thickness using methods previously described^16^. Brain regions (Figure 2) were punched or dissected out using a sterile scalpel in individual RNAase-free microcentrifuge tubes and stored at -80 until RNA extraction.

### RNA extraction and quantitative polymerase chain reaction (qPCR)

Brain tissue samples were removed from the -80ºC freezer and thawed on ice. Tissue was homogenized and RNA was purified and extracted using RNeasy Kit (Qiagen) following all manufacturer’s instructions. Isolated RNA samples were provided to the University of North Carolina Advanced Analytics Core to determine RNA concentration, synthesize and pre-amplify cDNA, and conduct high-throughput quantitative PCR (qPCR). qPCR was conducted using the Fluidigm Biomark HD 192.24 IFC array and validated mouse Taqman primers (ThermoFisher) according to the manufactures protocol. Relative gene expression was quantified using the 2^−ΔΔ^CT method, with target gene expression normalized to Gapdh and expressed relative to control samples. Samples (n=2) were excluded due to poor amplification or insufficient RNA concentrations.

### Statistical Analyses

Statistical analyses were conducted using SPSS v.29.0.1 (SPSS, Chicago, IL) and GraphPad Prism 10 (San Diego, CA). Differences in responses for operant conditioning and alcohol consumption were analyzed using a one-way ANOVA or two-way repeated measure ANOVAs where noted. Post hoc comparisons were performed using Tukey’s multiple comparisons test when a main interaction of group was identified. Group differences in gene expression were analyzed using 2 x 2 ANOVAs, with values reported as mean + SEM, and significance was defined as p < 0.05. A Spearman correlation was used to further explore associations between gene expression and both cumulative intake during binge drinking sessions and intake during operant conditioning. Significance testing was based on a bootstrapping procedure with 1000 samples and 95% CI.

## Results

We used the STAR framework to phenotype drinking behaviors. Mice were allowed 1-hr access to 15% v/v alcohol alone or adulterated with increasing concentrations of quinine, and responding was measured to evaluate individual consumption patterns. During this pre-binge operant testing, we did not see any differences in drinking patterns between the phenotypes (Figure 1B). Following this initial operant phase, mice were then given access to 15% v/v alcohol in a homecage using standard 2BC DiD procedures^6,11^. We did not see any significant drinking difference between groups during the first (Figure 1C) or second (Figure 1D) of homecage DiD access. When evaluating cumulative alcohol intake across both weeks of DiD, a one-way ANOVA revealed no significant differences between groups (Figure 1A; F(2, 21) = 2.093, P=0.1483). Following 2 weeks of DiD, mice were placed back into the operant chambers and given access to alcohol alone or adulterated with increasing concentrations of quinine in identical parameters and sessions as the pre-binge session. A two-way mixed-effects ANOVA showed a significant effect of phenotype F(2,21)=14.95, P<0.0001 and quinine concentration, F(2.433,51.10)=6.036, P=0.0026 and a significant phenotype x quinine concentration effect F(21,84)=1.804, P=0.0311 was observed (Figure 1F). A Tukey’s post hoc test revealed a significant difference between high (P<0.0001) and quinine-resistant (P=0.0399) vs low at the 0 μM concentration. In addition, a significant difference was noted between the quinine-resistant and low drinkers at the 250 (P=0.0019), 500 (P=0.0109), and 750 (P=0.0047) μM concentrations. Lastly, we also found significant differences between the high and quinine-resistant groups at the 750 (P=0.0005) and 1000 (P=0.0154) μM concentrations. Among the 24 mice, 9 (37.5%) mice were classified as low drinkers, 4 (16.7%) as high drinkers, and 11 (45.8%) as quinine resistant (Figure 1G). This distribution is comparable to that reported in the original study, which identified 46% low drinkers, 12% high drinkers, and 42% quinine-resistant drinkers. drinkers, and 42% as quinine-resistant drinkers^11^.

**Figure 1.**
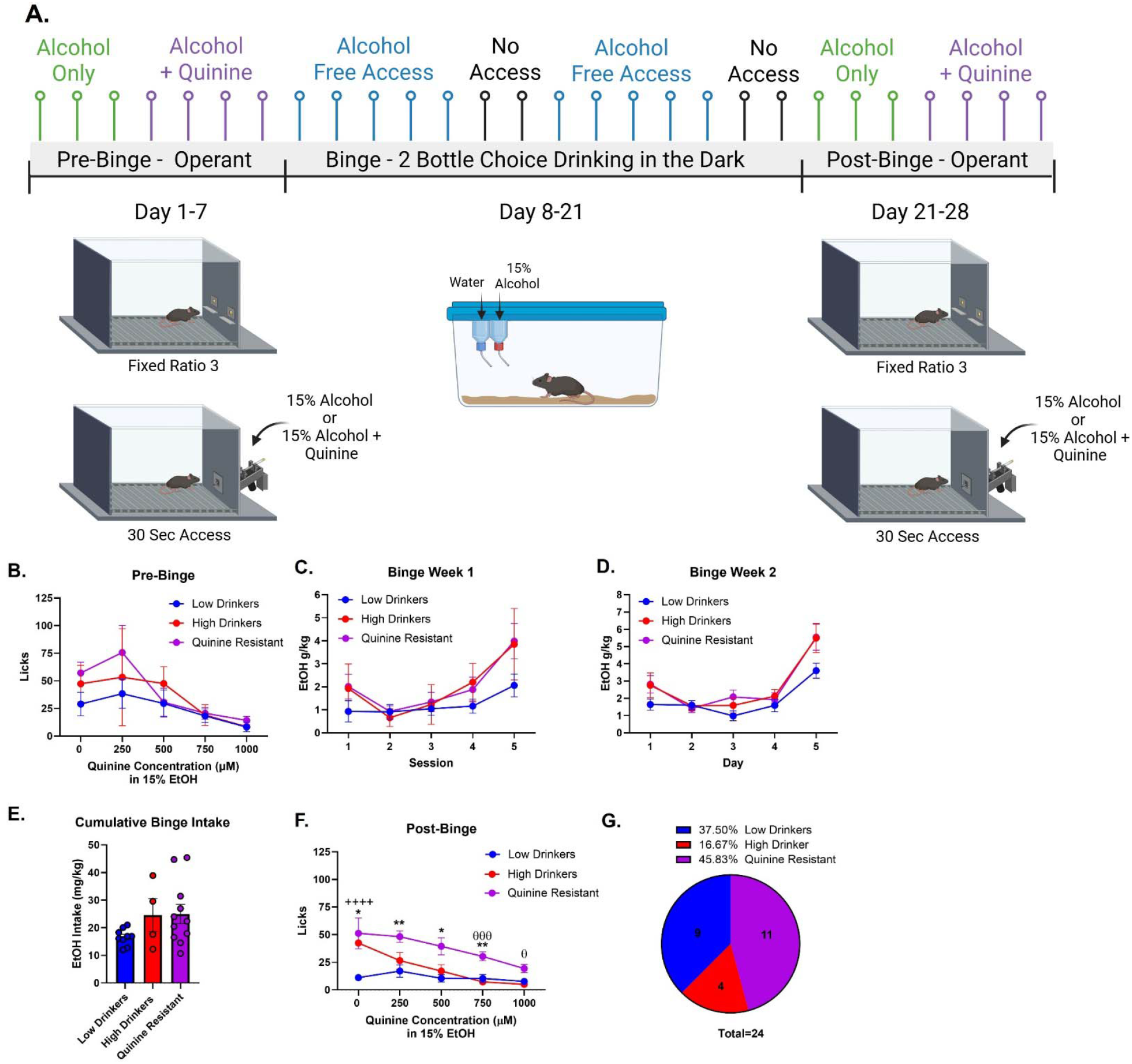
STAR methodology quantifies distinct alcohol drinking phenotypes. **(A)** STAR experimental design. Mice were exposed to alcohol under 3 phases: pre-binge operant responding, binge two bottle choice drinking in the dark, and a post-binge operant phase. **(B)** All mice consumed similar levels of 15% v/v alcohol alone (0 µM quinine) across and average of the 3 days of testing. They also consumed similar levels of quinine-adulterated alcohol across day 4-7. **(C)** Alcohol consumption during homecage 2BC DiD led to high levels of binge alcohol drinking during week 1 and week 2 **(D)**; however, no significant differences were notes between drinking phenotypes nor in cumulative intake across weeks **(E). (F)** During the post-binge phase, high and quinine resistant drinking consumed more alcohol alone on the average of day 1-3 of operant testing. When alcohol was adulterated with increasing concentrations of quinine, quinine resistant mice consumed more alcohol than low drinkers across all concentrations and high drinkers at the 750 and 1000 µM concentrations. Data are depicted as ± SEM. The following symbols represent: ^*^p<0.5, ^**^p<0.01 quinine resistant vs low drinkers; ^++++^p<0.0001 high drinkers vs low drinkers; ^θ^p<0.05, ^θθθ^p<0.05 high vs quinine resistant drinkers.

**Figure 2.**
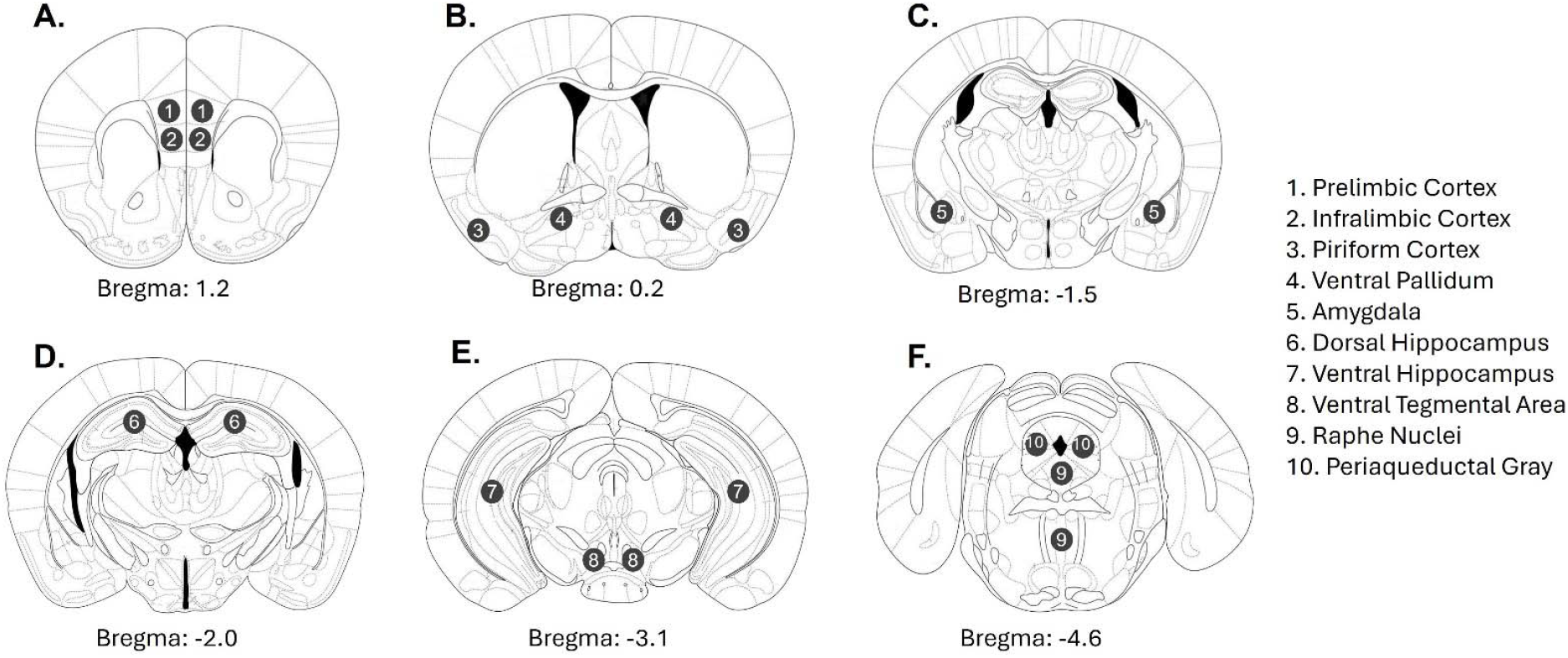
Following alcohol drinking, tissue was collected from 10 regions for qPCR analysis of gene expression alterations. Representative Tissue dissection diagram of punches for qPCR processing. All punches were collected frozen 1mm brain sections and regions were identified using landmarks from bregma coordinates 1.2 to -4.6. **(A)** From slices containing bregma AP 1.2, prelimbic (1) and infralimbic cortex was collected. **(B)** Piriform cortex (3) and ventral pallidum (4) was collected from slices containing bregma AP 0.2. **(C)** Amygdala (5) was collected from slices containing bregma AP -1.5 **(D)** Dorsal hippocampus (6) was collected from slices containing bregma AP -2.0. **(E)** Ventral Hippocampus (7) and ventral tegmental area (8) was collected from slices containing bregma AP - 3.1. **(F)** Raphe nuclei (9) and periaqueductal gray (10) was collected from slices containing bregma AP -4.6.

The day after the final operant session, mice were killed, and brain tissue was collected for qPCR. We first assess differences between low, high, and quinine-resistant drinkers with alcohol-naïve mice. We conducted One-way ANOVAs to assess group differences of the genes tested in Table 1. Complete statistical results of all comparisons are listed in Table 2, with all of the data included in an attached spreadsheet. For the following results, we have chosen to visualize only the comparisons that reached statistical significance or exhibited trends (P<0.1) towards significance.

**Table 1.** List of probes used for qPCR.

| Gene Name | Gene Abbreviation | Species | Assay ID |
| --- | --- | --- | --- |
| 5HT1A | Htr1a | Mouse | Mm00434106_s1 |
| 5HT1B | Htr1b | Mouse | Mm00439377_s1 |
| 5HT1D | Htr1d | Mouse | Mm00434115_s1 |
| 5HT1F | Htr1f | Mouse | Mm02619863_s1 |
| 5HT2A | Htr2a | Mouse | Mm00555764_m1 |
| 5HT2B | Htr2b | Mouse | Mm00434123_m1 |
| 5HT2C | Htr2c | Mouse | Mm00434127_m1 |
| 5HT3A | Htr3a | Mouse | Mm00442874_m1 |
| 5HT3B | Htr3b | Mouse | Mm00517424_m1 |
| 5HT4 | Htr4 | Mouse | Mm00434129_m1 |
| 5HT5A | Htr5a | Mouse | Mm00434132_m1 |
| 5HT5B | Htr5b | Mouse | Mm00439389_m1 |
| 5HT6 | Htr6 | Mouse | Mm00445320_m1 |
| 5HT7 | Htr7 | Mouse | Mm00434133_m1 |
| OCT-3 | Slc22a3 | Mouse | Mm00488294_m1 |
| Dopa Decarboxylase | DDC | Mouse | Mm00516688_m1 |
| Serotonin Transporter | Slc6a4 | Mouse | Mm00439391_m1 |
| Tryptophan Hydroxylase 2 | Tph2 | Mouse | Mm00557722_m1 |
| Kappa Opioid Receptor | Oprk1 | Mouse | Mm01230885_m1 |
| Nociceptin/orphanin FQ receptor | Oprl1 | Mouse | Mm00440563_m1 |
| Mu Opioid Receptor | Oprm1 | Mouse | Mm01188089_m1 |
| Beta Actin | ActB | Mouse | Mm00607939_s1 |
| Glyceraldehyde-3-phosphate dehydrogenase | Gapdh | Mouse | Mm99999915_g1 |

**Table 2.** Main effects of experimental groups on gene expression.

| Gene | Prelimbic Cortex | Infralimbic Cortex | Piriform Cortex | Ventral Pallidum | Amygdala | Dorsal Hipp. | Ventral Hipp. | VTA | PAG | Raphe |
| --- | --- | --- | --- | --- | --- | --- | --- | --- | --- | --- |
| Htr1a | F(3,29)=1.012, P=0.4016 | **F(3,30)=3.240, P=0.0358** | F(3,30)=1.088, P=0.3693 | F(3,29)=1.860, P=0.1585 | F(3,30)=1.422, P=0.2557 | F(3,30)=0.9348, P=0.4361 | F(3,30)=0.6616, P=0.5821 | F(3,30)=1.058, P=0.3817 | F(3,30)=0.7086, P=0.5544 | F(3,30)=0.5253, P=0.6683 |
| Htr1b | **F(3,29)=3.014, P=0.0460** | F(3,30)=2.897, P=0.0514 | F(3,30)=1.127, P=.539 | **F(3,29)=6.606, P=0.0015** | **F(3,30)=6.337, P=0.0019** | F(3,30)=2.605, P=0.0701 | F(3,30)=2.025, P=0.1315 | F(3,30)=0.1064, P=0.9557 | F(3,30)=2.164, P=0.1130 | F(3,30)=0.9224, P=0.4419 |
| Htr1d | **F(3,29)=3.040, P=0.0448** | F(3,30)=2.171, P=0.1121 | F(3,30)=0.6388, P=0.5960 | **F(3,29)=4.141, P=0.0147** | **F(3,30)=3.771, P=0.0208** | F(3,30)=1.209, P=0.3233 | F(3,30)=2.248, P=0.1031 | F(3,30)=0.3411, P=0.7957 | F(3,30)=2.344, P=0.0929 | F(3,30)=0.4994, P=0.6855 |
| Htr1f | F(3,29)=2.238, P=0.1050 | **F(3,30)=3.177, P=0.0383** | F(3,30)=0.4222, P=0.7384 | **F(3,29)=5.208, P=0.0053** | F(3,30)=1.935, P=0.1453 | F(3,30)=0.2793, P=0.8399 | **F(3,30)=3.259, P=0.0351** | F(3,30)=0.2116, P=0.8876 | F(3,30)=1.304, P=0.2912 | F(3,30)=0.1482, P=0.9300 |
| Htr2a | F(3,29)=1.101, P=0.3644 | F(3,30)=0.8183, P=0.4940 | F(3,30)=1.835, P=0.1620 | **F(3,29)=3.714, P=0.0225** | **F(3,30)=6.348, P=0.0018** | F(3,30)=1.338, P=0.2807 | F(3,30)=0.6241, P=0.6050 | F(3,30)=0.5171, P=0.6737 | F(3,30)=0.6805, P=0.5709 | F(3,30)=0.9751, P=0.4175 |
| Htr2c | F(3,29)=1.105, P=0.3631 | F(3,30)=1.322, P=0.2856 | F(3,30)=0.4118, P=0.7457 | F(3,29)=1.060, P=0.3814 | **F(3,30)=4.780, P=0.0077** | F(3,30)=2.326, P=0.0947 | F(3,30)=0.5525, P=0.6504 | **F(3,30)=3.081, P=0.0423** | F(3,30)=1.640, P=0.2010 | F(3,30)=0.4848, P=6954 |
| Htr3a | F(3,29)=0.4583, P=0.7135 | F(3,30)=2.532, P=0.0767 | F(3,30)=0.6199, P=0.6076 | F(3,29)=1.627, P=0.2047 | F(3,30)=0.4829, P=0.6967 | F(3,30)=1.792, P=0.1699 | F(3,30)=0.3274, P=0.8056 | F(3,30)=0.07434, P=0.9733 | F(3,30)=2.328, P=0.0945 | F(3,30)=0.6502, P=0.5890 |
| Htr4 | **F(3,29)=3.115, P=0.0414** | F(3,30)=0.3048, P=0.8217 | F(3,30)=0.5519, P=0.6508 | F(3,29)=2.411, P=0.0872 | F(3,30)=1.386, P=0.2661 | F(3,30)=1.039, P=0.3897 | F(3,30)=0.4649, P=0.7089 | F(3,30)=0.1907, P=0.9019 | F(3,30)=0.6819, P=0.5700 | F(3,30)=0.3931, P=0.7588 |
| Htr5a | F(3,29)=2.272, P=0.1012 | F(3,30)=1.619, P=0.2058 | F(3,30)=0.1601, P=0.9223 | F(3,29)=1.343, P=0.2796 | F(3,30)=2.027, P=0.1313 | F(3,30)=1.059, P=0.3812 | F(3,30)=0.8998, P=0.4528 | F(3,30)=1.018, P=0.3985 | F(3,30)=0.2500, P=0.8607 | F(3,30)=1.131, P=0.3524 |
| Htr5b | F(3,29)=0.4802, P=0.6986 | **F(3,30)=4.672, P=0.0086** | F(3,30)=1.068, P=0.3775 | F(3,29)=0.8437, P=0.4812 | F(3,30)=1.084, P=0.3708 | F(3,30)=0.9349, P=0.4360 | F(3,30)=0.9077, P=0.4490 | F(3,30)=0.07115, P=0.9749 | F(3,30)=0.2199, P=0.8818 | F(3,30)=1.837, P=0.1617 |
| Htr6 | F(3,29)=0.05338, P=0.6628 | F(3,30)=0.3714, P=0.7742 | F(3,30)=2.053, P=0.1275 | **F(3,29)=3.311, P=0.0338** | F(3,30)=2.212, P=0.1072 | F(3,30)=0.1791, P=0.9097 | F(3,30)=0.8878, P=0.4586 | F(3,30)=0.2935, P=0.8297 | F(3,29)=1.564, P=0.2194 | F(3,30)=0.3835, P=0.7656 |
| Htr7 | F(3,29)=0.2847, P=0.8360 | F(3,30)=2.370, P=0.0903 | F(3,30)=1.288, P=0.2966 | F(3,29)=1.286, P=0.2978 | F(3,30)=2.032, P=0.1305 | F(3,30)=0.2236, P=0.8792 | F(3,30)=0.6392, P=0.5957 | F(3,30)=1.461, P=0.2450 | F(3,30)=0.4717, P=0.7043 | F(3,30)=0.2359, P=0.8706 |
| Slc22a3 | F(3,29)=1.780, P=0.1730 | F(3,30)=0.9934, P=0.4093 | F(3,30)=0.8520, P=0.4765 | F(3,29)=0.2310, P=0.0971 | F(3,30)=2.002, P=0.1349 | F(3,29)=0.7893, P=0.3502 | F(3,30)=1.623, P=0.2049 | F(3,30)=0.1344, P=0.9388 | F(3,30)=0.1480, P=0.9301 | F(3,29)=2.721, P=0.0626 |
| DDC | F(3,29)=1.417, P=0.2577 | F(3,30)=0.6137, P=0.6115 | F(3,30)=0.6181, P=0.6087 | F(3,29)=2.141, P=0.1166 | F(3,30)=2.2760, P=0.0594 | **F(3,30)=3.029, P=0.0447** | F(3,30)=1.281, P=0.2988 | F(3,30)=0.2341, P=0.8719 | F(3,30)=0.3354, P=0.7998 | **F(3,30)=3.056, P=0.0434** |
| Slc6a4 | - | - | - | - | - | - | - | - | - | F(3,28)=1.730, P=0.1835 |
| Tph2 | F(3,29)=0.4794, P=0.6991 | F(3,30)=1.027, P=0.3948 | F(3,30)=0.6817, P=0.5702 | F(3,29)=0.3656, P=0.7783 | F(3,30)=0.8001, P=0.5036 | F(3,30)=0.7512, P=0.5303 | F(3,30)=0.4243, P=0.7370 | F(3,30)=1.328, P=0.2836 | F(3,30)=0.1077, P=0.9550 | **F(3,30)=3.497, P=0.0275** |
| Oprk1 | F(3,29)=0.08498, P=0.9677 | F(3,30)=0.4828, P=0.6968 | F(3,30)=2.622, P=0.0688 | **F(3,29)=3.282, P=0.0348** | **F(3,30)=6.991, P=0.0011** | F(3,30)=0.4916, P=0.6908 | F(3,30)=0.6531, P=0.5873 | F(3,30)=0.8587, P=0.4732 | F(3,30)=0.9423, P=0.4325 | F(3,30)=0.5871, P=0.6280 |
| Oprl1 | F(3,29)=0.1156, P=0.9502 | **F(3,30)=3.236, P=0.0360** | F(3,30)=0.8205, P=0.4928 | F(3,29)=1.107, P=0.3623 | **F(3,30)=3.072, P=0.0427** | F(3,30)=0.8160, P=0.3127 | **F(3,30)=5.187, P=0.0052** | **F(3,30)=3.591, P=0.0250** | F(3,30)=0.4813, P=0.6978 | F(3,30)=0.2862, P=0.8349 |
| Oprm1 | F(3,29)=2.443, P=0.0841 | F(3,30)=1.772, P=0.1737 | F(3,30)=0.5665, P=0.6413 | F(3,29)=2.147, P=0.1158 | **F(3,30)=4.066, P=0.0155** | F(3,30)=1.546, P=0.2222 | F(3,30)=0.9945, P=0.4088 | F(3,30)=0.9396, P=0.4338 | F(3,30)=0.5928, P=0.6246 | F(3,30)=0.7588, P=0.5260 |

In the prelimbic cortex, we found significant main effects of the 5-HT1B (Figure 3A), 5-HT1D (Figure 3B), 5-HT4 (Figure 3C), genes, and a trend towards significance for the OPRM1 gene (Figure 3D). Posthoc analyses revealed that low drinkers had significant increases in the expression of 5-HT1B (Figure 3A, P=0.0452) and 5-HT1D (Figure 3B, P=0.0420) compared to control mice. In addition, we found that quinine-resistant drinkers displayed increased expression of 5-HT1B (Figure 3A, P=0.0338) and 5-HT4 (Figure 3C, P=0.0338) compared to controls. Drinking experience led to increased expression of 5-HT1A (Figure 3E), 5-HT1F (Figure 3G), 5-HT5B (Figure 3I), and trends towards altered expression of 5-HT1B (Figure 3F), 5-HT3A (Figure 3H), and 5-HT7 (Figure 3J). Posthoc analysis showed quinine-resistant mice had increased expression of 5-HT1A (Figure 3E, P=0.0216), 5-HT1F (Figure 3G, P=0.0128). Expression of 5-HT5B (Figure 3I) was found to be increased in both high drinkers (P=0.0478) and quinine-resistant drinkers (P=0.0039) compared to controls.

**Figure 3.**
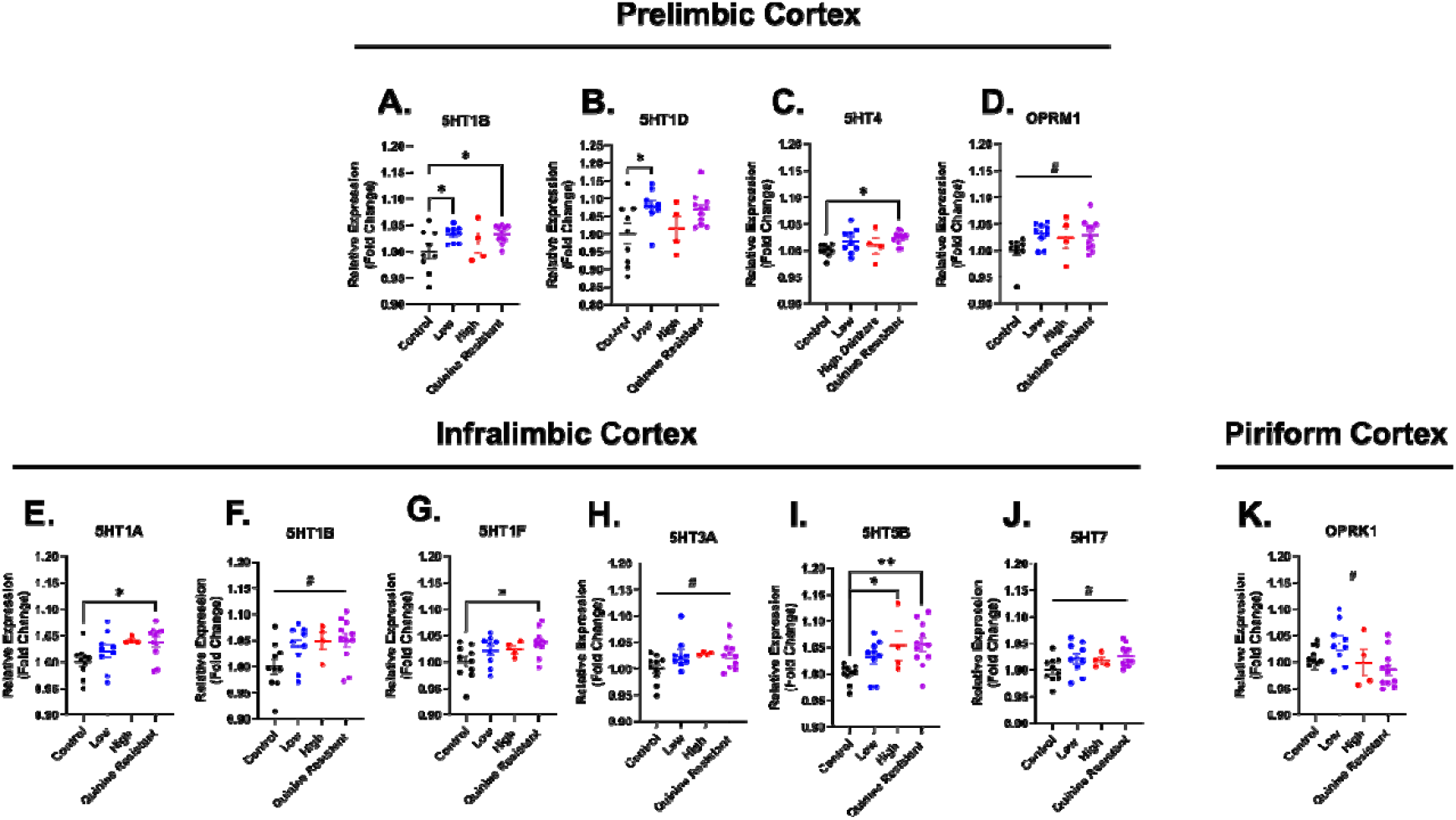
Chronic alcohol drinking led to alterations in gene expression in prelimbic, infralimbic, and piriform cortex in a phenotype specific manner. In the prelimbic cortex, greater expression of 5HT1B **(A)**, 5HT1D **(B)**, 5HT4 **(C)** was higher in alcohol drinking mice and a trend towards significance in OPRM1 **(D)**. In the infralimbic cortex, greater expression of 5HT1A **(E)**, 5HT1F **(G)**, 5HT5B **(I)** was seen in alcohol consuming mice compared to controls. A trend towards significance was noted in 5HT1B **(F)**, 5HT3A **(H)** and 5HT7 **(J)**. In the piriform cortex, only a trend was noted in OPRK1 **(K)**. Data are depicted as ± SEM. ^*^p<0.5, ^**^p<0.01. #=trending; p<0.01.

Across all regions assessed, the ventral pallidum and amygdala demonstrated the greatest extent of gene differences amongst groups. In the ventral pallidum we found that chronic alcohol drinking led to increased expression of 5-HT1B (Figure 4), 5-HT1D (Figure 4B), 5-HT1F (Figure 4C), 5-HT2A (Figure 4D), 5-HT6 (Figure 4F), and OPRK1 (Figure 4H). Expression of 5-HT4 (Figure 4E) and OCT3 (Figure 4G) showed a trends towards significance in differing expression between the groups. Quinine-resistant mice showed a significant increase in expression of 5-HT1B (Figure 4, P=0.0007), 5-HT1D (Figure 4B, P=0.0057), 5-HT1F (Figure 4C, P=0.0027), 5-HT2A (Figure 4D, P=0.0089), 5-HT6 (Figure 4F, P=0.0487), and OPRK1 (Figure 4H, P=0.0264) as revealed by post-hoc analyses. Low drinkers also displayed increased expression of 5-HT1B (Figure 4A, P=0.0239), 5-HT1F (Figure 4C, P=0.0347), and 5-HT6 (Figure 4F, P=0.0375) compared to alcohol-naïve controls. Similar to ventral pallidum, the amygdala showed a similar increase in expression of 5-HT1B (Figure 4I), 5-HT1D (Figure 4J), 5-HT2A (Figure 4K), and OPRK1 (Figure 4N) in drinking-experienced animals compared to controls. We also found altered expression of 5-HT2C (Figure 4L), OPRL1 (Figure 4O), OPRM1 (Figure 3P), and a trend towards significance for DDC (Figure 4M) between groups. Interestingly, in contrast to the effects seen in the ventral pallidum, high drinking animals showed the greatest increase in expression of many of the genes including in the amygdala. Posthoc analyses revealed a significant increase in expression of 5-HT1B (Figure 4I, P=0.0007), 5-HT1D (Figure 4J, P=0.0118), 5-HT2A (Figure 4K, P=0.0013), 5-HT2C (Figure 4L, P=0.0151), OPRK1 (Figure 4N, P=0.0003), OPRL1 (Figure 4O, P=0.0410), and OPRM1 (Figure 3P, P=0.0055). We also found increased expression of 5-HT2A expression in quinine-resistant mice (Figure 4K, P=0.0080) and 5-HT1B expression in low and quinine-resistant mice (Figure 4I, P=0.0449 and P=0.0464, respectively) compared to controls.

**Figure 4.**
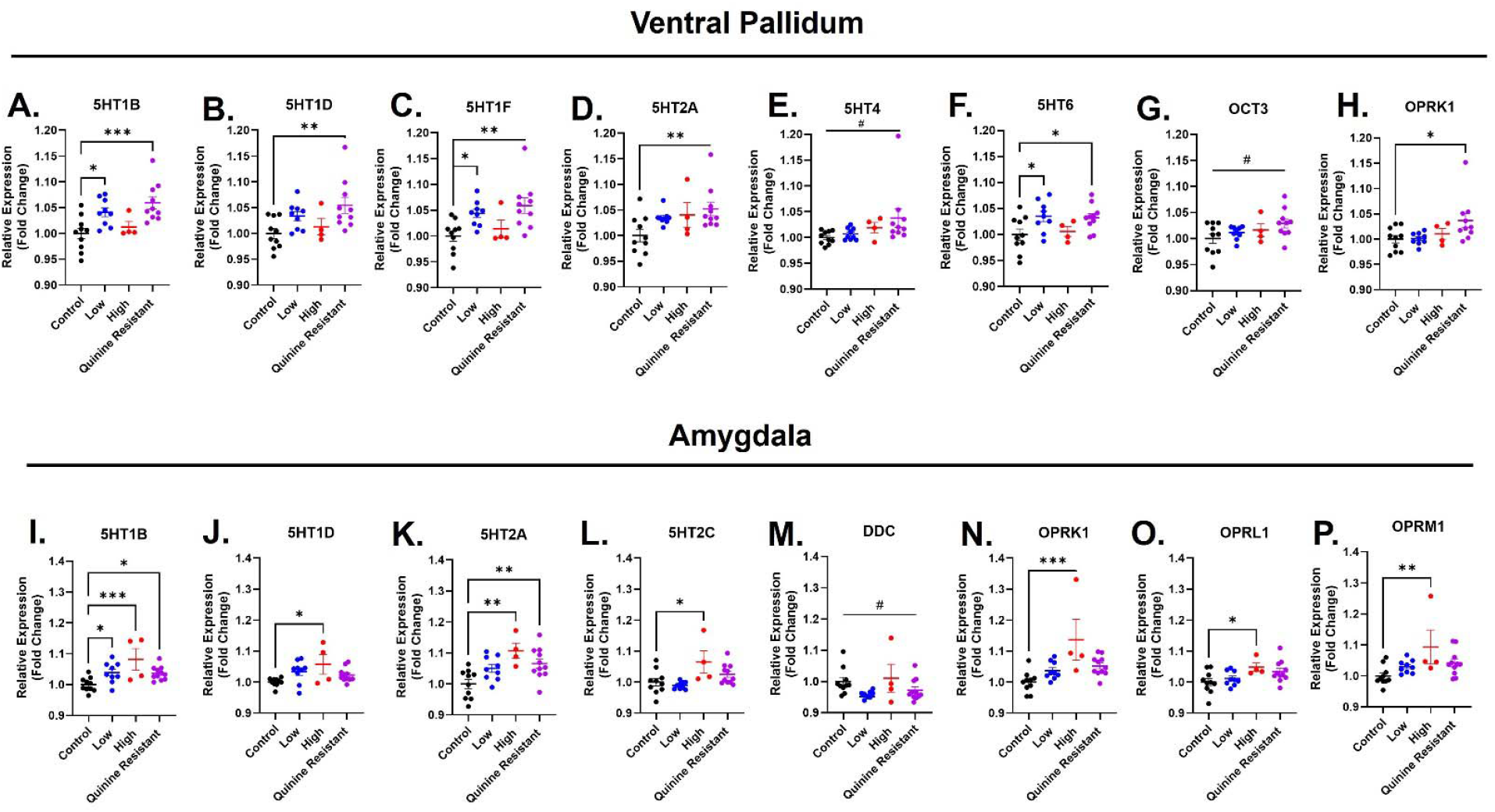
Alcohol drinking led to altered gene expression in the ventral pallidum and amygdala in a phenotype-dependent manner. Analysis of the ventral pallidum revealed greater expression of 5HT1B **(A)**, 5HT1D **(B)**, 5HT1F **(C)**, 5HT2A **(D)**, 5HT6 **(F)**, and OPRK1 **(H)** was higher in alcohol drinking mice. Expression of 5HT4 **(E)** and OCT3 **(G)** was trending towards a difference between groups. In the amygdala, greater expression of 5HT1B **(I)**, 5HT1D **(J)**, 5HT2A **(K)**, 5HT2C **(L)**, OPRK1 **(N)**, OPRL1 **(O)**, and OPRM1 **(P)** was seen in alcohol consuming mice compared to controls. A trend towards significance was noted in DDC **(M)**. Data are depicted as ± SEM. ^*^p<0.5, ^**^p<0.01, ^***^p<0.001. #=trending; p<0.01.

Finally, in mid-to hind-brain regions (Figure 5) we saw a different pattern of gene alteration than the aforementioned regions. In the dorsal hippocampus we found opposing direction of gene alterations between groups with 5-HT1B (Figure 5A) being increased in the alcohol consuming mice and 5-HT2C (Figure 5B) displaying a trend towards decreased expression. Posthoc analyses found a significant increase in 5-HT1B expression in low drinkers (Figure 5A, P=0.0262) compared to controls. We also found a significant reduction in DDC (Figure 5K) expression with posthoc tests revealing trends in significance for all groups compared to control. In the ventral hippocampus we found significant increases in expression of 5-HT1F and OPRL1 with posthoc analyses showing significant increases in 5-HT1F expression in low drinkers (Figure 5L, P=0.0205) and OPRL1 expression in high (Figure 5M, P=0.0026) and quinine-resistant (Figure 5M, P=0.0308) drinkers compared to control mice. The VTA display increased expression of 5-HT2C and OPRL1 alcohol drinking mice. Posthoc analyses found a trend towards significance in expression of 5-HT2C in quinine-resistant mice (Figure 5A, P=0.1032) and significant increase in expression of OPRL1 in quinine-resistant (Figure 5B, P=0.0416) and high drinking (Figure 5B, P=0.0291) mice compared to controls. Alcohol drinking led to a reduction in DDC and TPH2 expression in the raphe nuclei. We also found a trend towards significant reduction in expression OCT3. Posthoc analyses showed the expression of DDC was significantly reduced in low and quinine-resistant mice compared to control (Figure 5H, P=0.0472 and P=0.0298, respectively) as well as TPH2 (Figure 5J, P=0.0319 and P=0.0214, respectively). Lastly, the PAG showed trends of altered expression of 5-HT1D and 5-HT3A only.

**Figure 5.**
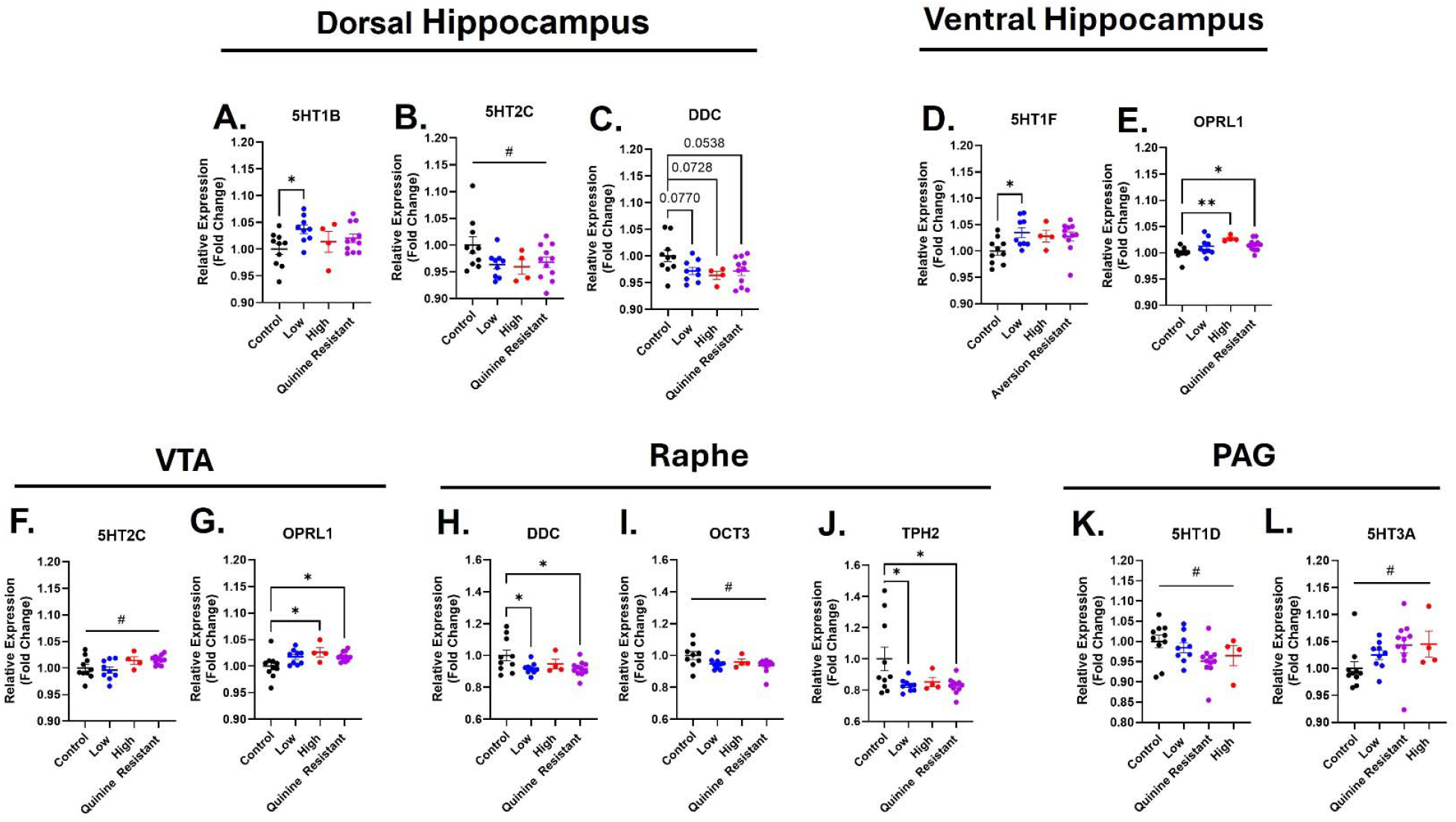
Alcohol drinking led to altered gene expression in the hippocampal and midbrain regions in a phenotype-dependent manner. Alcohol drinking mice displayed increased expression of 5HT1A **(A)** but reduced expression of DDC **(C)** in the dorsal hippocampus. Expression of 5HT2C (B) was trending towards significance. In the ventral hippocampus, expression of 5HT1F **(D)** and OPRL1 **(E)** was higher in alcohol drinking mice. The VTA revealed altered expression of OPRL1 **(G)** and a trend towards significance in 5HT2C expression **(F)**. In the raphe nuclei, DDC **(H)** and TPH2 **(J)** expression was reduced in alcohol drinking mice. A trend towards a significant reduction in the expression of OCT3 **(I)** in alcohol drinking mice compared to controls was noted. The PAG revealed only trends towards significantly altered expression of 5HT1D **(K)** and 5HT3A **(L)**. Data are depicted as ± SEM. ^*^p<0.5, ^**^p<0.01, ^***^p<0.001. #=trending; p<0.01.

We next evaluated the relationship between alcohol intake during the post-binge (phenotyping) phase and gene expression changes by correlating cumulative alcohol consumption across the three alcohol-only operant sessions (Figure 1) to identify markers that are responsive to high alcohol intake. Bootstrapped correlation analyses were conducted to determine significant relationships between alcohol intake and gene expression. For clarity only correlations that remain significant after bootstrapping sampling are shown. We found that alcohol consumption during post-binge phase positively correlated with 5-HT5A in the prelimbic region (Figure 6A, Spearman’s ρ = -0.445, 95% CI [-0.728, -0.054], P=0.029). We also noted significant positive relationship between the ventral pallidum expression of 5-HT1A (Figure 6B, Spearman’s ρ = -0.419, 95% CI [0.015, 0.711], P=0.041), 5-HT5A (Figure 6B, Spearman’s ρ = 0.476, 95% CI [0.138, 0.743], P=0.019), and 5-HT7 (Figure 6C, Spearman’s ρ = 0.525, 95% CI [0.185, 0.797], P=0.008) and alcohol alone consumption. Conversely, expression of DDC in the dorsal hippocampus showed a negative correlation with operant alcohol consumption (Figure 6D, Spearman’s ρ = -0.443, 95% CI [-0.680, -0.092], P=0.029).

**Figure 6.**
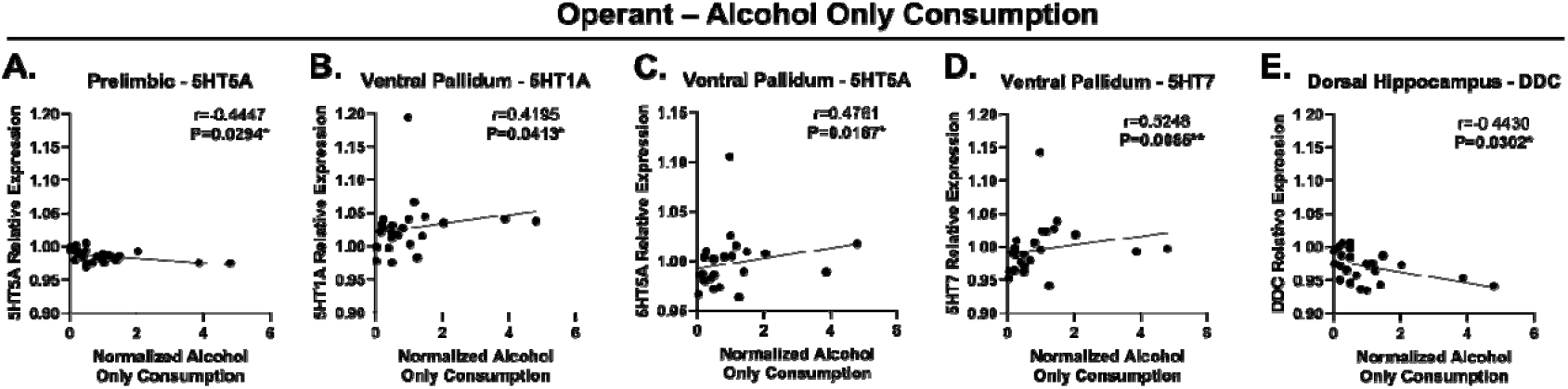
Alcohol drinking during the post-binge operant phase was correlated with altered gene expression in the prelimbic cortex, ventral pallidum, and dorsal hippocampus. **(A)** Expression of 5HT5A in the prelimbic cortex was negative correlated with 15% v/v alcohol (unadulterated) drinking. In the ventral pallidum, expression of 5HT1A **(B)**, 5HT5A **(C)**, and 5HT7 **(D)** was positively correlated with alcohol only consumption. **(E)** In the dorsal hippocampus, alcohol drinking was found to be negatively correlated with expression of DDC. ^*^p<0.5, ^**^p<0.01 r = Spearman correlation coefficient.

We hypothesized that aversion-resistant drinking would engage distinct neural mechanisms; thus, we examined gene expression changes that correlated with increasing quinine-adulterated intake during the post-binge operant session. Two regions emerged as having significantly altered expression that correlated with increasing quinine-adulterated drinking: the ventral pallidum and the PAG. Within the ventral pallidum, we found that quinine-adulterated drinking positively correlated with the expression of 5-HT1A (Figure 6A, Spearman’s ρ = 0.528, 95% CI [0.110, 0.826], P=0.008), 5-HT1B (Figure 6B, Spearman’s ρ = 0.510, 95% CI [0.148, 0.765], P=0.011), 5-HT5A (Figure 6C, Spearman’s ρ = 0.540, 95% CI [0.162, 0.147], P=0.006), 5-HT7 (Figure 6D, Spearman’s ρ = 0.523, 95% CI [0.088, 0.760], P=0.009), and OPRK1 (Figure 6E, Spearman’s ρ = 0.564, 95% CI [0.196, 0.794], P=0.004). In the PAG, we found a significant positive correlation between quinine-adulterated drinking and expression of 5-HT2C (Figure 6F, Spearman’s ρ = 0.537, 95% CI [0.130, 0.230], P=0.007), 5-HT3A (Figure 6G, Spearman’s ρ = 0.570, 95% CI [0.260, 0.810], P=0.004), and OPRL1 (Figure 6G, Spearman’s ρ = 0.446, 95% CI [0.065, 0.738], P=0.029).

While we evaluated the relationship between altered expression and non-adulterated alcohol consumption during the operant sessions, we questioned whether consumption during the 2BC DiD sessions may lead to distinct changes in 5-HT and opioid machinery. During the 2BC DiD sessions we hypothesized this was the phase where the mice received the bulk of their alcohol exposure as the daily exposure time was longer (2 or 4 hr for DiD sessions vs 1hr for operant sessions) and the alcohol drinking was non contingent. Therefore, we compared cumulative alcohol intake for 10 sessions of 2BC DiD with gene expressions. Surprisingly, we only found that two genes correlated with 2BC DiD intake. Expression of 5-HT1F in the amygdala (Figure 7A, Spearman’s ρ = -0.483, 95% CI [-0.820, -0.001], P=0.017) and 5-HT3A in the ventral hippocampus (Figure 7B, Spearman’s ρ = - 0.509, 95% CI [-0.792, -0.087], P=0.011) revealed a significant negative correlation with cumulative alcohol intake during this phase.

**Figure 7.**
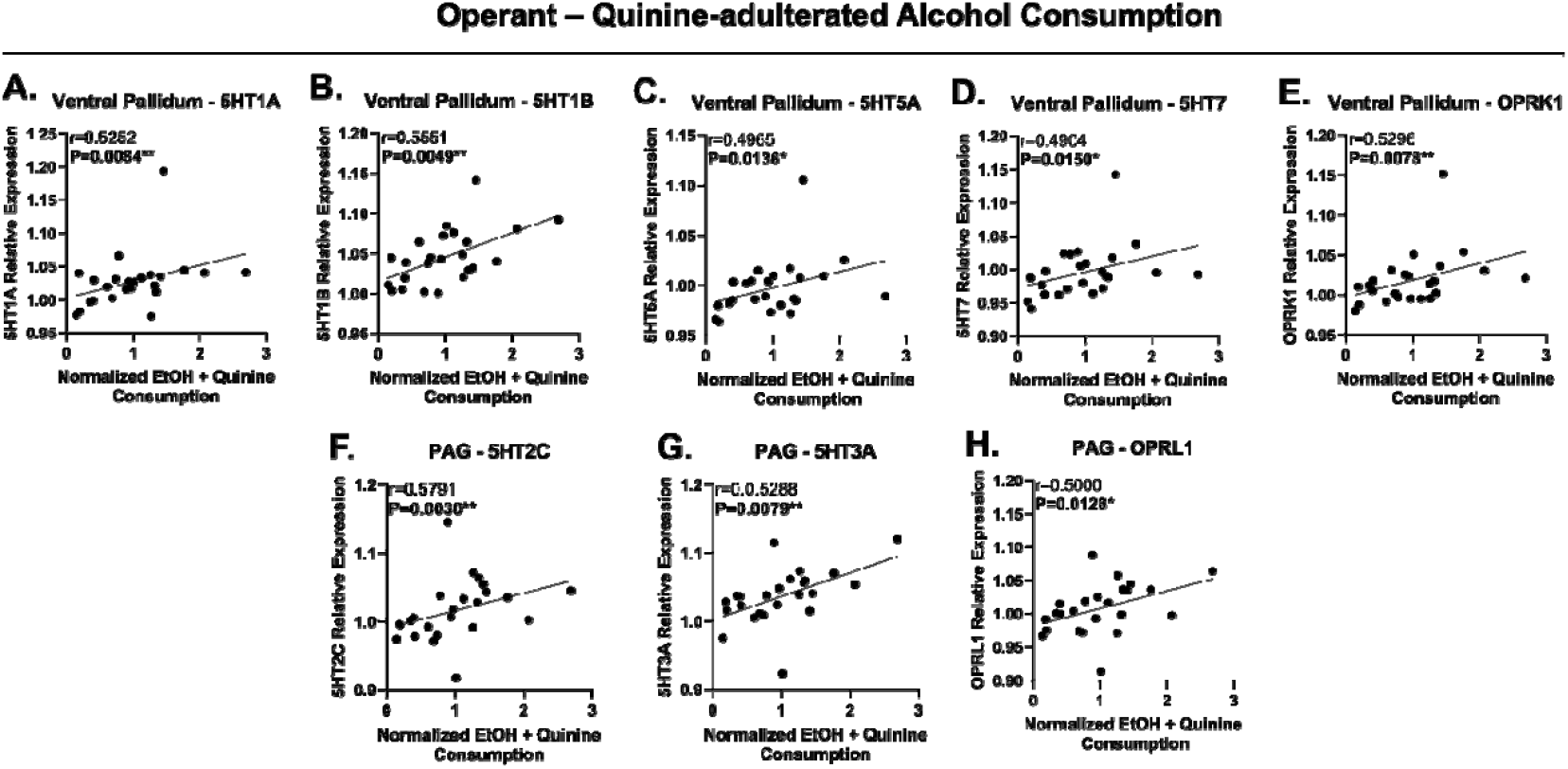
Quinine-adulterated alcohol drinking during the post-binge operant phase was significantly positively correlated with expression of 5HT and opioid markers in the ventral pallidum and periaqueductal gray. Expression of 5HT1A **(A)**, 5HT1B **(B)**, 5HT5A **(C)**, 5HT7 **(D)**, and OPRK1 **(E)** in the ventral pallidum was positively correlated with cumulative quinine adulterated drinking across all 4 concentrations (250, 500, 750, 1000 µM). In the PAG, expressions of 5HT2C **(F)**, 5HT3A **(G)**, and OPRL1 **(H)** were positively correlated with quinine adulterated drinking. ^*^p<0.5, ^**^p<0.01 r = Spearman correlation coefficient.

**Figure 8.**
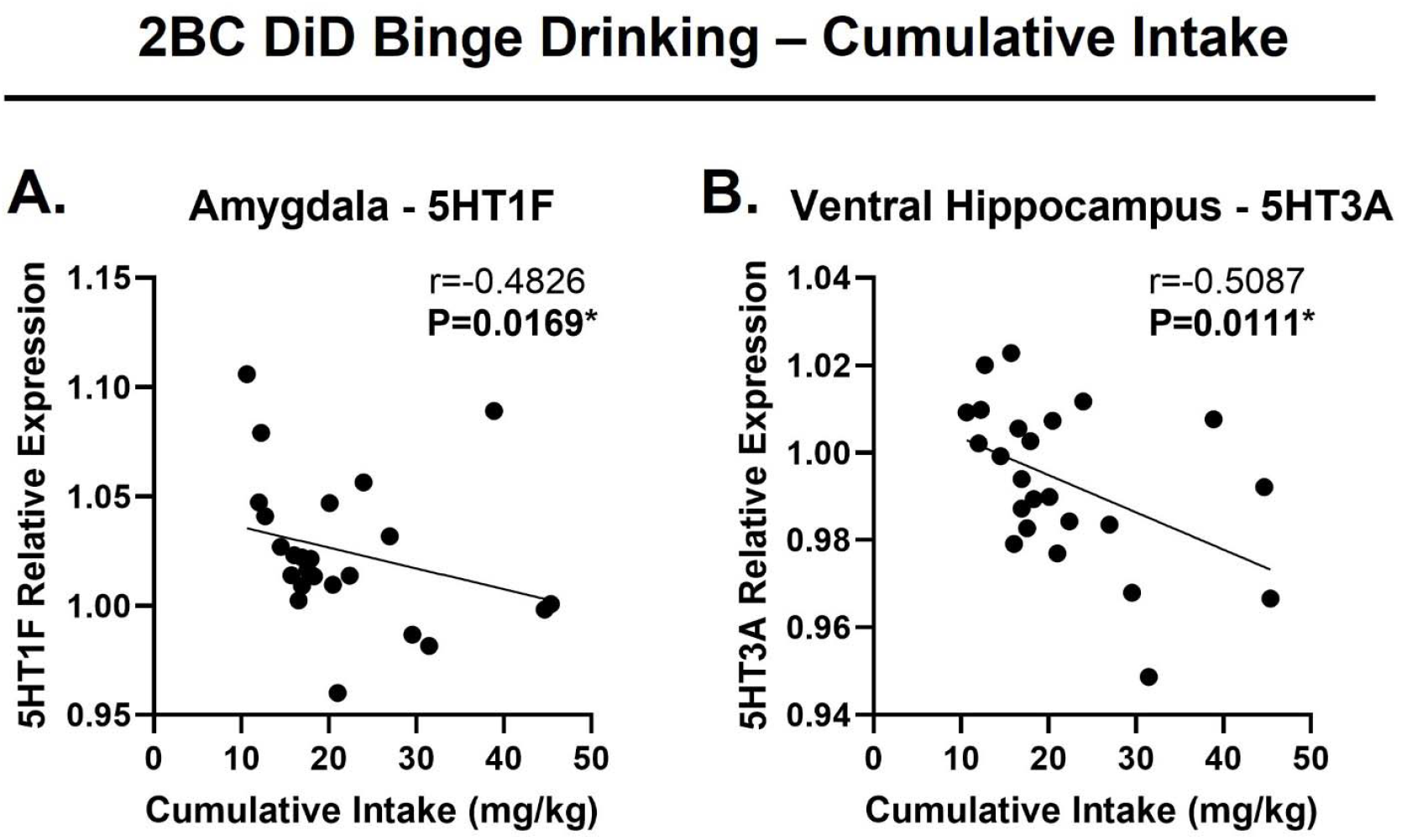
Binge drinking during homecage 2BC DiD was negatively associated with expression of 5HT receptors in the amygdala and ventral hippocampus. **In the amygdala**, Expression of 5HT1F **(A)** was negatively correlated with cumulative alcohol intake during the 2 weeks of DiD. Expression of 5HT3A in the ventral hippocampus was also negative correlated with cumulative alcohol intake during the 2 weeks of DiD. ^*^p<0.5, ^**^p<0.01 r = Spearman correlation coefficient.

## Discussion

The goal of the present study was to describe adaptations in theserotonin and opioid system function that may drive problem alcohol use vulnerability. Here, we identify region-specific alterations in the mRNA for anumber of 5-HT GPCRs, transporters, and synthesis enzymes in addition to changes in opioid GPCRs relative to alcohol-naïve control mice. Further, we assessed the relationship of these altered genes with endophenotypes of AUD, including high intake of alcohol and aversion-resistant drinking and found significant molecular correlates of these behavioral patterns. Collectively, these findings extend prior work implicating serotonergic and opioid systems in alcohol use by identifying novel, region-specific molecular markers that are associated with AUD-relevant drinking endophenotypes.

Serotonin containing cell bodies are located in the hindbrain nuclei, specifically the DRN and MRN. Acutely, alcohol leads to a reduction in firing rates of raphe nuclei^17–19^ and increased extracellular levels in downstream regions such as the prefrontal cortex^20^, striatum^17,21^, amygdala, and hippocampus^18^. Chronic exposure to the physiological properties of raphe nuclei has been mixed, with some reporting decreased firing^18^ and others reporting increased firing^22^. However, studies have shown that chronic drinking has led to reduced 5-HT levels^23^, metabolites^12^, and transporter binding and expression^24^. Together, suggesting that chronic alcohol leads to reduced 5-HT neurotransmission. Our findings support and expand these findings in that we found reductions in DDC, an enzyme critical for the final catalytic step of 5-HT synthesis, and TPH2, the enzyme encoding the rate-limiting step for synthesis. When assessing how these markers relate to the drinking metrics measured, we found that expression of the 5-HT2C and 5-HT3A receptors in the PAG was positively correlated with quinine-adulterated drinking. Recently, a study found that 5-HT2C and 5-HT3A receptors in the dorsal PAG interact to regulate anxiety-like behavior,^25^ further supporting that serotonin regulation plays a critical role in mediating reward and aversive properties. Our findings suggest serotonin functionis reduced in the raphe cell bodies after alcohol drinking. Reduced 5-HT function after chronic drinking would lead to changes in downstream signaling in fore- and mid-brain regions. One way these adaptations can occur is by up- or downregulating receptors to change inhibitory or excitatory drive-in attempt to restore homeostasis as shown in the current study. Below we will expand on findings in these downstream regions in light of previous literature.

Chronic drinking led to an increased expression of G_i/o_-protein coupled 5-HT receptors across both the prelimbic and infralimbic cortices. This is supported by a study that demonstrated elevated 5-HT1A density^26^ and binding sites^27^ in the frontal cortex of alcohol preferring (P) rats. Another study, showed chronic drinking in non-human primates also led to increased 5-HT1A binding in frontal cortex subregions^28^. Many of the 5-HT1 receptor subtypes, including 5-HT1A, 5-HT1B, and 5-HT1D, function as both somatodendritic and terminal autoreceptors, respectively, as well as postsynaptic heteroreceptors^29^. Here, we measured mRNA expression, and therefore the results likely reflect changes in postsynaptic heteroreceptor expression as presynaptic machinery would be located in the cell bodies in raphe nuclei, though mRNA levels don’t necessarily predict protein abundance or functional output^30^. In the medial prefrontal cortex (including both prelimbic and infralimbic cortices), 5-HT1 receptors are located on excitatory pyramidal neurons, GABAergic interneurons, and parvalbumin interneurons^31^. Given findings demonstrating increased inhibitory tone in the prefrontal cortex after chronic alcohol^32,33^, it’s possible that the if upregulated expression as indicated in the current study is a result of increased expression on pyramidal neurons, leading to reduced excitatory drive and shifting the balance towards increased inhibitory tone. Future studies investigating a more direct role of 5-HT1 receptor function in modulating neuronal subtypes of the prefrontal cortex are warranted. In addition to 5-HT1, we found that expression of the 5-HT5A receptor in the prelimbic cortex significantly correlates with the amount of alcohol consumed during the post-binge operant phase. To our knowledge, the role of 5-HT5A receptors in the prelimbic cortex remains to be explored. However, there is evidence that these receptors are expressed throughout layer II and IV of the cortex^34^ and have a functional role in regulating the synaptic plasticity of the prelimbic cortex^35^. Furthermore, 5-HT5A has been implicated in sleep disruptions, cognitive function, and mood disorders (for review see^36^). Together, this suggests that the alcohol-induced reductions seen in the expression of inhibitory regulators in the current study may drive changes in top-down control of these neurological functions through regulation of the excitability of the cortex.

Of note, our findings demonstrate that a majority of the 5-HT and opioid gene alterations after alcohol were found in the ventral pallidum and amygdala regions. While the amygdala and its associated nuclei have long been recognized as central regulators of alcohol-related behaviors and as a key hub for serotonergic modulation of affective, arousal, and motivational processes, the ventral pallidum has only recently begun to receive comparable attention^37–39^. Here, we found that alcohol led to elevated gene expression of 5-HT serotonin receptors of the 5-HT1, 2, 4, and 6 family in the ventral pallidum, as well as 5-HT1 and 5-HT2 receptors in the amygdala. The ventral pallidum receives moderately dense input from the dorsal raphe nucleus^40–42^ and possesses many of the 5-HT receptor subtypes, including from the 5-HT-1, -2, -3, and -4 classes^43–48^. One study has shown that tissue content of ventral pallidum punches had nearly 2-fold greater levels of 5-HT and the serotonin metabolite, 5-hydroxyindoleacetic acid (5-HIAA), than either dopamine or its metabolite, 3,4-dihydroxyphenylacetic acid (DOPAC). However, the large majority of alcohol and drug-related studies in the ventral pallidum have focused on dopaminergic signaling (for review see^37,38,49^). We found that alcohol led to upregulation of expression of many of the serotonin receptors, and this increased expression was consistently higher in the quinine-resistant mice. Further, expression of these receptors was found to be tightly correlated with alcohol drinking behavior, regardless of whether presented alone or quinine adulterated. Given the critical role of 5-HT in mediating drug and alcohol relapse seeking^50^ and aversive behaviors^51^, it’s likely that the increased expression of serotonin receptors could be a factor driving alcohol-related behaviors. This is supported by a recent study showing that cocaine-induced inhibition of GABAergic synaptic transmission in the ventral pallidum is mediated by serotonin transporter blockade, resulting in increased synaptic serotonin and subsequent activation of 5-HT1B receptors, rather than canonical dopamine signaling^48^. We also found increased mRNA expression of the kappa opioid receptor (KOR) in quinine-resistant mice, and that expression was positively correlated with increased consumption of quinine-adulterated alcohol. These effects are in contrast to one study that found microinfusion of KOR agonists or antagonists in the ventral pallidum had no effect on voluntary homecage alcohol drinking in alcohol preferring rats^52^; however, they did not assess aversion-resistant drinking. Collectively, these findings suggest that serotonergic and KOR signaling may play discrete roles in regulating aversion-resistant alcohol drinking and underscore the need for mechanistic studies examining serotonergic function within the ventral pallidum in reward- and aversion-related alcohol behaviors.

Similar to the ventral pallidum, in the amygdala we found increased expression of 5-HT1 and 5-HT2 receptors as well as all 3 opioid receptors in alcohol consuming mice; though, in contrast, increased expression was seen predominantly in the high alcohol drinking phenotype. Our finding that alcohol drinking led to increased 5-HT1B expression in all drinking phenotypes and 5-HT1D in high drinkers is interesting in light of other studies suggesting that systemic administration of 5-HT1B agonists leads to a reduction in alcohol drinking^53,54^, and alcohol preferring rats have reduced 5-HT1B density in the amygdala^55^. However, because these approaches, including the current findings, do not isolate individual amygdala nuclei or cell populations, the specific contribution of 5-HT1B signaling remains unclear. Autoradiography mapping has allowed for the identification of the 5-HT1D receptor^56–59^ in the amygdala of human and rodent brains; yet the role and function of this receptor is unknown. Additionally, we found that 5-HT1F expression positively correlated with cumulative intake during DID. Considering the greatest upregulation of 5-HT1 receptors was seen in high drinking mice and that this class of receptor relates to intake during a binge drinking procedure, it’s likely that this class of receptors in the amygdala plays a critical role in mediating excessive alcohol drinking. In contrast, substantially greater attention has been devoted to the 5-HT2 class in studies of amygdala and alcohol-related behaviors. Studies have documented increased 5-HT2C binding density in the amygdala of alcohol preferring rats^60^. Other studies found that microinfusion of a non-selective 5-HT2 agonist in the basolateral amygdala attenuated alcohol seeking but had only moderate effects on consumption of a sweetened alcohol solution^61^, and alcohol dependence and withdrawal disrupted central amygdala 5-HT2C signaling without altering expression^62^. Here, we found elevated expression of 5-HT2A in all drinking phenotypes and 5-HT2C in high drinking phenotypes only. Given that antagonism of 5-HT2C in the amygdala reduced alcohol withdrawal-related anxiety measures^63,64^ and the strong link between alcohol use and affective dysregulation, 5-HT2C is a likely mechanism central to this relationship, as proposed by others^65^.

Another critical mechanism regulating the relationship between alcohol drinking and affective disorders is the opioid system. We found that high drinkers displayed upregulated mRNA expression of mu, kappa, and nociception opioid receptors in the amygdala. These findings are well supported by a host of studies showing that antagonism of these receptors in the amygdala reduces alcohol consumption, seeking, and relapse vulnerability among other alcohol-related behaviors^66–72^. Additionally, clinical studies report increased binding of mu and nociception opioid receptors^73,74^ in alcohol dependent subjects; however, reduced kappa receptor availability in the amygdala of alcohol dependent subjects was also reported^75^. Thus, a better understanding of nucleus-specific serotonergic and opioid signaling within the amygdala will be critical for clarifying how these systems regulate alcohol-related behaviors.

Within the dorsal hippocampus, we found that alcohol consumption led to increased 5-HT1B expression and a trend towards reduction in expression of 5-HT2C, suggesting a shift in the excitatory-to-inhibitory (E/I) balance as others have demonstrated^76^. One study showed that chronic drinking in non-human primates led to a reduction in serotonin transporter binding in the CA1 of the hippocampus^77^, suggesting that there may be a compensatory downregulation due to low serotonin levels. In another study, this group also reported that chronic drinking in monkeys led to increased expression of 5-HT1A binding in the CA1 but no change in mRNA expression^78^. Our findings mirror many of these in that we found increased expression of another 5-HT1 receptor, 5-HT1B, and reduction of DDC, a critical regulator of the conversion of 5-hydroxytrptophan (5-HTP) to serotonin, suggesting reduced serotonin levels and upregulated 5-HT1 receptors. Furthermore, we found this change in DDC expression was significantly negatively correlated with alcohol alone consumed in operant chambers, suggesting that excessive drinking leads to a reduction in synaptic serotonin levels in the dorsal hippocampus. This is consistent with a study showing chronic alcohol and spontaneous withdrawal resulted in supersensitized 5-HT1A receptors and reduced 5-HT synthesis in the hippocampus of rats^79^. We evaluated the dorsal and ventral hippocampus separately because they are canonically implicated in distinct functional domains, with the dorsal hippocampus primarily involved in learning and memory and the ventral hippocampus more closely regulating affective and emotional behaviors^80^. Results from the ventral hippocampus revealed increased 5-HT1F and nociception opioid receptor expression in alcohol drinking mice. While no studies have investigated the role of ventral hippocampus 5-HT1F receptors in AUD, their expression has been confirmed^81^. Though due to being inhibitory coupled, it’s possible the alteration of these receptors may alter hippocampal excitability following chronic alcohol exposure as they’ve been found to be colocalized with glutamate neurons in the brain^82^. The NOP receptor is highly expressed in the hippocampus^83^, and alcohol preferring rats express higher basal levels of nociception/orphanin FQ (N/OFQ) in the hippocampus^84^. However, studies have shown that chronic alcohol leads to reduced = pronociceptin expression in humans^85^ and N/OFQ expression in the hippocampus of rodents^86^. Though we didn’t find any significant correlations with OPRL1 expression and the behavioral metrics evaluated, it’s possible that these findings more closely reflect expression conferring vulnerability to alcohol consumption rather than alcohol-induced adaptation. Along these lines, our findings did demonstrate that 5-HT3A expression was negatively correlated with cumulative intake during binge drinking in the dark session. 5-HT3 receptors are highly expressed on GABAergic interneurons,^87^ where stimulation of these ligand-gated ion-channels leads to increased GABA release^88^. Our results suggest that chronic, excessive alcohol may lead to a compensatory downregulation of ventral hippocampus 5-HT3 expression, thereby reducing GABA levels and shifting towards excitatory drive^76^. Together, these findings suggest a role in which compensatory changes in serotonin function may shift E/I ratios and alter intrinsic excitability within the hippocampus. Future studies investigating cell-type specific expressions of these changes will help distinguish whether these effects reflect alcohol-induced adaptations or pre-existing vulnerability factors.

The VTA serves as a central locus for the mesolimbic dopaminergic system and has long-standing implications in the alcohol field. It also receives dense serotonin projections from raphe nuclei, suggesting serotonin may play a critical role in regulating the dopaminergic pathway. Here, we found that alcohol drinking mice had increased 5-HT2C and NOP opioid receptor expression compared to alcohol-naïve controls. Given that serotonin is known to regulate dopamine firing through activation of 5-HT2C on GABAergic interneurons and thereby inhibit dopamine neurons in the VTA^89,90^, increased expression would suggest that alcohol’s ability to disinhibit dopamine neurons would be reduced. Additionally, alcohol-elicited dopamine release downstream in the nucleus accumbens would also be dampened, requiring increased alcohol consumption to reach comparable dopamine levels. This occurrence is supported in the 2C expression in low drinkers, which was identical to control and only elevated in high alcohol consuming and quinine-resistant phenotypes. Future work is required to elucidate serotonergic control of VTA dopamine circuits and its role in alcohol-related behaviors. We also identified an upregulation in NOP mRNA expression in alcohol drinking mice. Microinfusion of a

NOP receptor antagonist in the VTA reduces alcohol drinking under 2BC without altering water or food intake in alcohol preferring rats^91^. Together, these findings suggest heightening NOP signaling and/or expression following chronic alcohol exposure, and that NOP antagonists may serve as a potential pharmacotherapeutic for excessive alcohol drinking.

In sum, our results identify region-specific serotonergic and opioid markers that may mediate alcohol drinking phenotype or be altered following chronic alcohol exposure. One limitation of these findings is that we are not able to identify which markers suggest a predisposed vulnerability to heavy alcohol drinking or if they are markers of alcohol-induced adaptations due to chronic exposure. Additionally the current study only used male mice. In the initial STAR study, females exhibited phenotypic responses comparable to males when assessed using the same parameters, despite lower alcohol consumption during the operant phases, suggesting that this framework is applicable to both sexes. A recent review article noted the lack of research on serotonin function in females and emphasized the need for further studies^92^, therefore future studies in females are warranted. However, by identifying these behaviorally-relevant biomarkers, we have begun unraveling the interplay of serotonin and opioids functioning in AUD pathophysiology. Further research into the gaps in scientific literature identified here will aid in producing new pharmacotherapeutics and treatments of AUD.

## Notes

### Competing Interest Statement

The authors have declared no competing interest.

### Summary of Updates

I mis-spelled an authors name, and left a note about word limit in abstract.

### Funding

- NIH: AA019454, AA031604

